# Benthic diversity along an Arctic fjord: which are the key factors?

**DOI:** 10.1101/2023.08.16.553514

**Authors:** Anaïs Lebrun, Steeve Comeau, Frédéric Gazeau, Jean-Pierre Gattuso

## Abstract

Arctic coastal ecosystems include benthic communities that hold an important role within the marine food chain. Kelps, fucoid species, and coralline algae dominate rocky habitats, offering food and shelter for various species. Kelps and fucoid species also aid in carbon sequestration, sediment stabilization, and erosion mitigation. In summer, the influx of freshwater from glacier and permafrost melt alters coastal waters conditions. The input of turbid freshwater influences underwater light, salinity, and substrate, impacting benthic organism distribution. This study investigates possible link between environmental conditions and benthic diversity through environmental DNA (eDNA). Six sites were monitored along Kongsfjorden (Svalbard,Norway) during the summer of 2021. Contrary to expectations, macroalgal distribution didn’t correlate with light, and suspension feeders showed no clear links with chlorophyll a or nutrient concentrations. Glacial influence may have contributed to higher benthic diversity. Predators’ presence, tied to glacier proximity, possibly explained this trend. However, further studies are needed to validate these observations and assumptions.

## Introduction

Arctic coasts host a fragile ecosystem characterized by diverse benthic communities that play a crucial role in supporting the entire marine food web. In rocky areas, benthic communities are dominated by kelps, fucoid species, and coralline algae (Lebrun et al., 2022). These species provide food and habitat for fish, crabs, and many other species. Kelps and fucoid species play important roles in carbon sequestration and help stabilize sediments and reduce coastal erosion (Løvås and Tørum, 2001; Filbee-Dexter et al., 2019). Soft-substrate communities are generally less diverse and present lower biomass but represent around 65% of the Arctic coastal areas due to the river and glacial runoffs and are dominated by polychaetes and specialized malacostracans (Lantuit et al., 2012; Filbee-Dexter et al., 2019). In both substrates, the benthic fauna such as mollusks, echinoderms, and arthropods is essential in recycling organic matter and maintaining ecosystem health (Welsh et al., 2003; März et al., 2021).

In the summer months, melting glaciers and permafrost discharge large amounts of freshwater to Arctic coastal areas. As glaciers melt, they release sediment that has been trapped in the ice. Furthermore, as the meltwater from land-terminating glaciers and permafrost flows toward the fjord, it gets enriched with sediment (Hopwood et al., 2018). As a result, when meltwater reaches the fjord, it carries a significant sediment load. The input of turbid freshwater affects the local underwater light regime, salinity, nutrients, and substrate, and, hence, the distribution and abundance of benthic organisms. From the inner to the outer part, Arctic fjords are characterized by a gradient of substrate ranging from soft to harder and gradient. Also, in summer the fjord displays a distinct a gradient of underwater light and salinity with turbid and low saline waters in the inner parts to less turbid and more saline ones in the outer parts (Svendsen et al. 2002; Kedra et al., 2010). The benthic diversity is largely influenced by the substrate type and the water characteristics such as turbidity and nutrient content (Anderson et al., 2008, Saeedi et al., 2022).

With climate change, the Arctic is experiencing rapid environmental changes with temperatures rising at more than twice the global average rate (Richter-Menge et al., 2017). As temperatures rise, glacier melting is accelerating and occurs earlier in the year. Erosion of permafrost in coastal areas in the Arctic increased by 80 to 160%, in comparison to average rates from the ‘80s (Jones et al., 2020). This leads to an intense discharge of freshwater and sediment into the fjord (Howe et al., 2010). The increase in runoff expands the area impacted by glacier melting and therefore impacts organisms which, until now, were not affected by turbid and fresh waters (Barnhart et al., 2016).

In this study, we investigated how the benthic flora and fauna are distributed along an Arctic fjord. The main objective was to characterize potential variations in community composition related to changes in environmental conditions. We selected six sites along the Kongsfjorden (Svalbard, Norway) and monitored their environmental parameters (temperature, salinity, photosynthetically active radiation (PAR), nutrient, chlorophyll *a* (chl *a*) concentration, pH_T,_ and total alkalinity) and benthic diversity during the summer of 2021. Diversity was assessed using environmental DNA (eDNA), a molecular tool allowing taxonomic identification from part of an organism’s DNA, found in its environment. Our hypothesis is that the distribution of benthic organisms is impacted by the gradient of light, salinity, and substrate type. In particular, we hypothesize that the distribution of macroalgae follows the light gradient and that of suspension feeders follows the salinity gradient, associated with the chlorophyll *a* (chl *a*) gradient.

## 1. Material and methods

### 1.1. Sampling sites

Kongsfjorden is an open fjord situated on the north-western coast of Svalbard (12°E, 79°N). It hosts four tidewater glaciers (Kronebreen, Kongsbreen, Conwaybreen, and Bloomstrandbreen) on its north and east edges. We monitored the environmental parameters and diversity at six sites along the fjord from June to August 2021 (Kongsfjordneset 78.97°N, 11.48°E; Hansneset 78.99°N, 11.98°E; Bloomstrand East 78.97°N, 12.19°E; Ossian Sars 78.92°N, 12.45°E; French Bird Cliff 78.90°N, 12.20°E; and Kongsbreen South 78.91°N, 12.54°E; Fig. 1)

**Figure 1:**
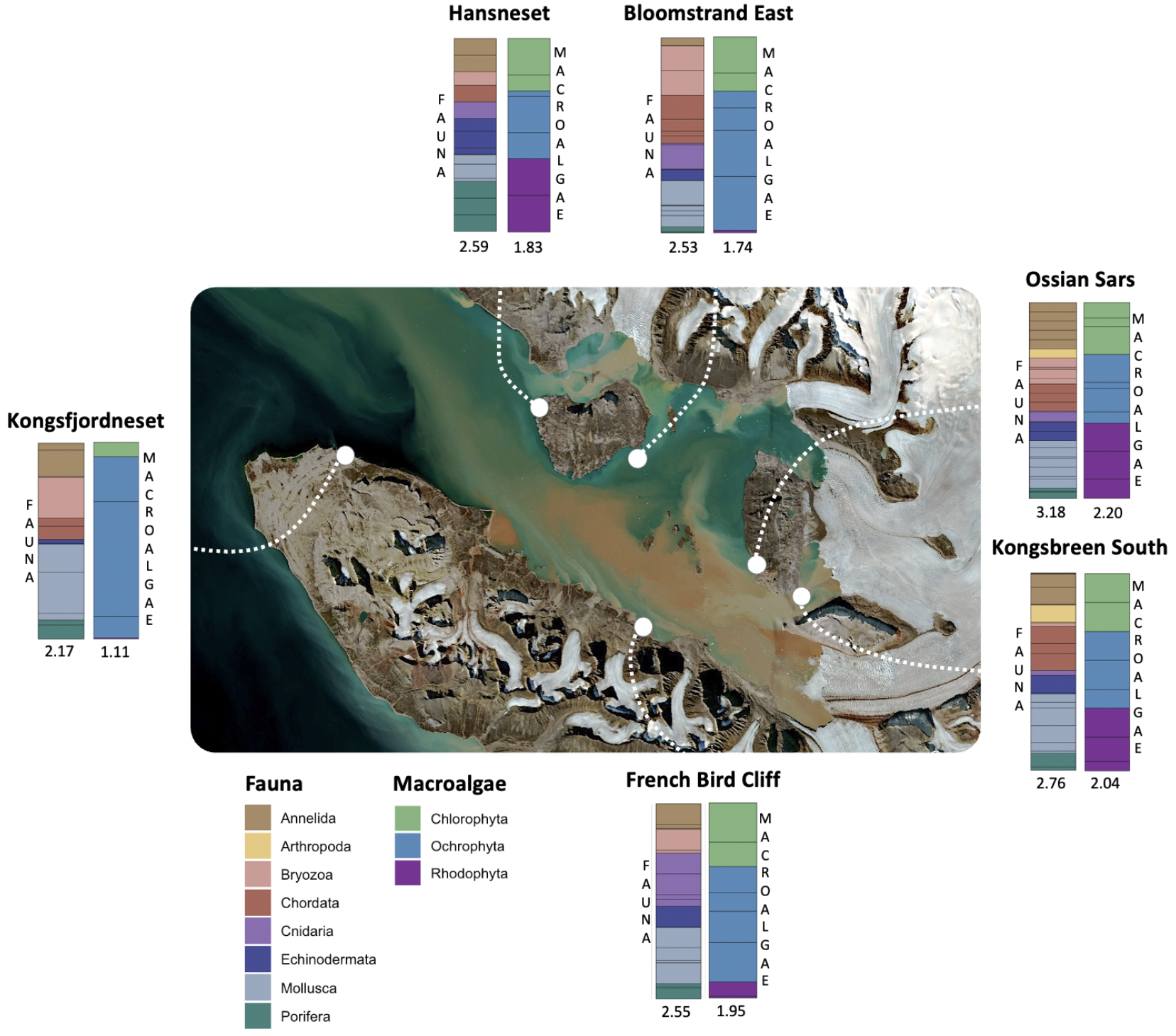
Satellite image of Kongsfjorden (Sentinel 2 L2A, 2020/07/27, available on sentinel-hub.com consulted on 2023/01) and the six sampling locations with stacked barplot representing eDNA faunal and macroalgal diversity at the order level. Phyla are represented by color. The size of the cells represents the proportion of the orders (separation lines) and phyla (colors) based on the eDNA indexes. The values of the Shannon diversity index are indicated below each barplot.

### 1.2. Environmental parameters

At the six sites, we weekly sampled seawater for nutrient concentration (NO_2_, NO_3_, PO_4_, SIOH_4_), total alkalinity, pH_T_ (on the total scale), and chl *a* concentration using a 12 L Niskin bottle closed 1 meter above the bottom. Salinity and temperature profiles were performed using a CTD (Seabird SBE-19). PAR profiles were conducted using a planar light sensor connected to a data logger (LICOR LI-400) to determine the vertical light attenuation coefficient (K_d_) by linear regression between depth and the log-transformed irradiance. Three sampling times for environmental parameters (2021-06-24 & 26, 2021-07-22 & 23, and 2021-08-26 & 27) coincided with the eDNA samplings (see section 3.1.).

#### 1.2.1. Nutrient concentration

Seawater samples for nutrient concentration measurements were filtered (porosity: 0.2 μm), transferred to acid-washed 60 mL polyethylene vials, and stored in the dark at − 20 °C until analysis. All nutrient samples were analysed using a standard automated colorimetry system (Seal Analytical continuous flow AutoAnalyser III, AA3) at the Institut de la Mer de Villefranche (IMEV, France). Nitrate (NO_3_) and nitrite (NO_2_) ions were analysed according to the protocol of Bendschneider and Robinson (1952) with a detection limit of 0.05 μM and 0.01 mM, respectively. The concentration of phosphate (PO_4_) ions was determined following the protocol developed by Murphy and Riley (1962), with a detection limit of 0.02 μM. Silicate (SIOH_4_) ions were analysed according to the protocol of Strickland and Parsons (1972) with a detection limit of 0.05 μM and 0.02 mM.

#### 1.2.2. pH_T_ and total alkalinity

Seawater samples for pH_T_ measurements were collected directly from the Niskin in 300 mL borosilicate bottles with a stopper and were analyzed the same day. pH_T_ (pH on the total scale) was determined by measuring absorbances at 434, 578, and 730 nm on a Cary60 UV spectrophotometer (Agilent) before and after the addition of 50 μL of purified meta-cresol purple (R. H. Byrne, University of South. Florida) as described by Dickson et al. (2007). pH_T_ was calculated using formulae and constants described by Liu et al. (2011).

A volume of 500 mL of seawater was filtered on GF/F membranes for total alkalinity (*A*_T_) measurements. *A*_T_ was determined potentiometrically using a Metrohm© titrator (Titrando 888) and a glass electrode (Metrohm©, Ecotrode Plus) calibrated using NBS buffers (pH 4.0 and pH 7.0, to check that the slope was Nernstian) and a TRIS buffer solution (salinity 35; provided by Andrew Dickson, Scripps University, USA). The accuracy of the data was validated by titrating a standard seawater provided by Andrew Dickson (Scripps University, USA). Duplicate titrations were performed on 50 mL sub- samples and *A*_T_ was calculated as described by Dickson et al. (2007).

#### 1.2.3. Chl *a* concentration

A volume of 500 mL of seawater samples was filtered on GF/F membranes for chl *a* concentration measurements and stored at −80 °C. Chl *a* was extracted in 90% aqueous acetone for 24 h in the dark at 4 °C. After cold-centrifugation (0 °C, 15 min, 3000 rpm), the supernatant was transferred into glass vials and F_0_, the initial fluorescence of chl *a* and pheophytin pigment, was measured using a fluorimeter (Turner Design 10-AU Fluorimeter). The F_a_ fluorescence was measured one min after the addition of 10 μl of 0.3 N chlorhydric acid to transform chl *a* into pheophytin pigment and subtract Fa from Fo. The concentration of chl *a* was calculated as described by Lorenzen (1967).

### 1.3. eDNA

#### 1.3.1. Sampling and extraction

We sampled monthly (06-24 & 26, 07-22 & 23, and 08-26 & 27) 1 L of water in triplicate at each of the six sites using a 12 L Niskin bottle that was closed 1 meter above the substrate at 6 to 8 m depth. Back at the Kings Bay Marine Laboratory, samples were filtered through cellulose acetate filters (25 mm, porosity: 0.45 μm, Sartorius) using a peristaltic pump (WATSON MARLOW) and stored at −80 °C. DNA collected on filters was extracted using 200 μL of QuickExtract DNA extraction solution (Lucigen, Illumina) and 0.4 g of beads (glass beads 0.75A 1.0 mm, Dutscher) to dislodge the genetic material from the filter and put it in solution. After 4 min of vortex, the samples were incubated two times (6 min at 65 °C and 2 minutes at 98 °C) with 1 min of vortex between each step, before being centrifuged (8000 rpm, 30 sec). The supernatants were stored at −20 °C. DNA extracts were purified from potential PCR inhibitors using the ReliaPrep DNA Clean-up kit (Promega).

#### 1.3.2. PCR amplification and sequencing

The DNA extracts were amplified using a PCR amplification reaction targeting 1) a 313 bp fragment encoding cytochrome c oxidase I (COI, Leray et al., 2013) and 2) the V4 SSU 18S region of 381 bp encoding 18S ribosomal DNA (18S; Stoeck et al., 2010). These gene regions allow the taxonomic identification of eukaryotes. We used the Multiplex MasterMix (Qiagen) and followed the PCR touchdown program of Leray et al. (2013) for both primers. High-throughput sequencing was performed by the Génome Quebec platform (Canada) using an Illumina MiSeq.

#### 1.3.3. Bioinformatics

The primers were trimmed using cutadapt (version 4.2) and bash scripts written by Ramon Gallego (Kelly lab). Demultiplexing and matching of forward and reverse reads were performed using the dada2 software package (version 1.26.0) and script written by Elizabeth Andruszkiewicz Allan (Kelly lab, github.com/KellyLabUW/MiSeqPipeline). Sample replicates that did not meet the required quality standards were excluded from the study (25 out of 106 for COI primer and 21 out of 106 for 18S primer). The remaining biological and technical replicates were merged together by sites and sampling period to run the data analysis. The BLASTn (nucleotide-nucleotide Basic Local Alignment Search Tool) was performed using GenBank (NCBI) with the Octopus server from LBDV (Laboratoire de Biologie du Développement, Villefranche-sur-Mer).

The number of reads was normalized to the eDNA index (Kelly et al., 2019) to minimize amplification biases between taxa. Only benthic taxa found in two of the three sampling periods were kept for analysis to ensure reliable and consistent data as well as temporal stability.

### 1.4. Statistics

Multiple Hutcheson t-tests were used to test the significance of the difference between communities’ Shannon diversity indexes (package ecolTest). Correlation tests were used to test for the link between biodiversity and environmental parameters.

## 2. Results

### 2.1. Environmental characterization of the sites

The environmental data at each site was used to establish a grouping by site and/or period (Fig. 2). Data are shown in the supplementary material (Fig S1). When considering all the measured parameters, the Kongsbreen South site from week 28 to 34 (early July to the end of August) and the Ossian Sars site on week 33 clearly differed from the other sites with high nutrient concentrations and a high K_d_ (Fig. 2A). With a high salinity and total alkalinity, weeks 27 and 28 at French bird cliff and 32 at Kongsfjordneset also differed from the rest of the sites.

**Figure 2:**
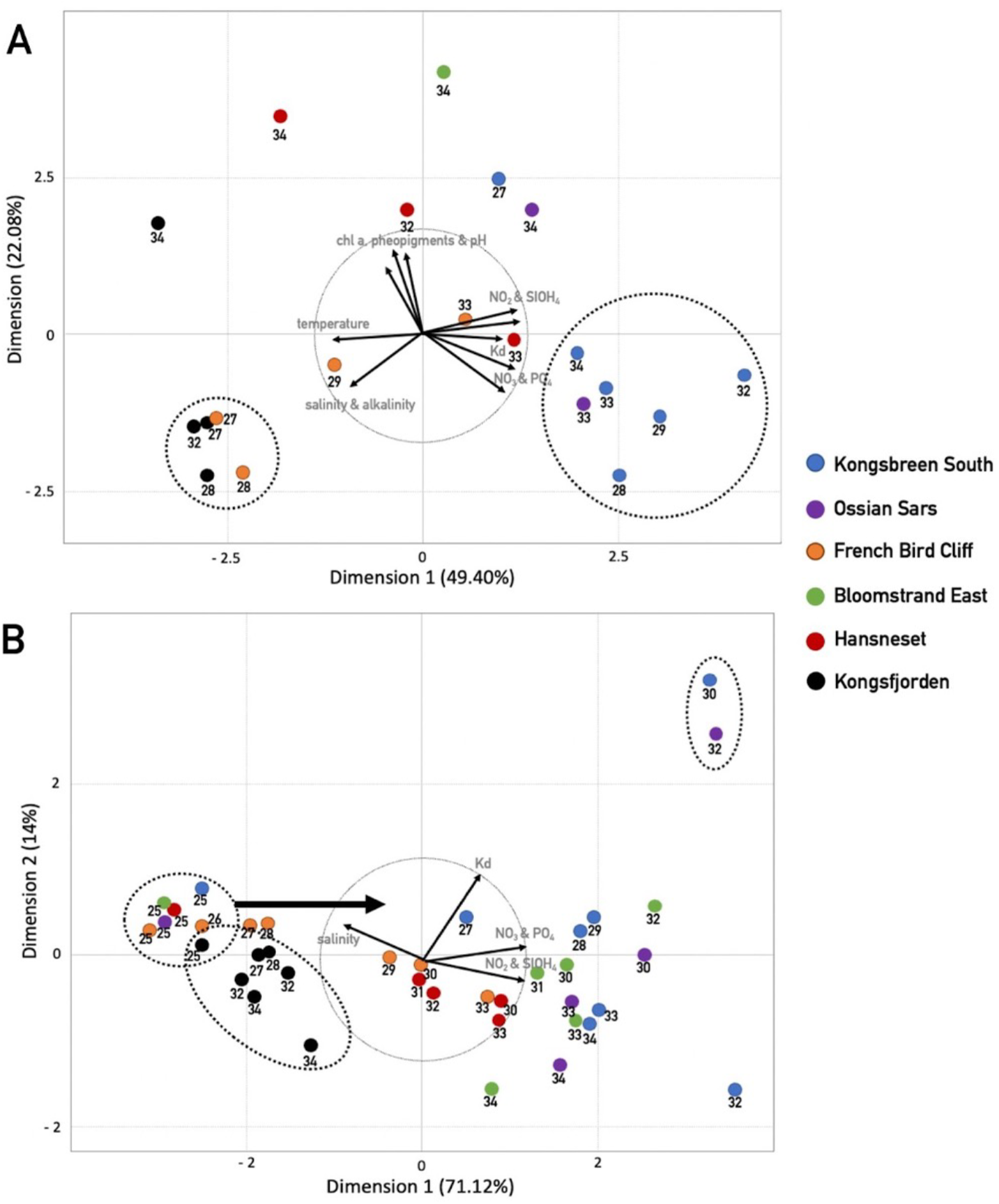
Principal Component Analysis (PCA) based on A) all parameters or B) salinity, K_d_, and nutrient concentrations measured from week 25 to week 34 at six study sites along the Kongsfjorden. The numbers on the figure indicate the week of sampling. The bold arrow in B) represents the movement of sites from left to right of the graph after week 25.

Temperature and K_d_ exhibited an inverse relationship because turbid waters are located in the innermost part of the fjord where temperatures are at their lowest. K_d_ and nutrient concentrations were positively correlated because turbid meltwaters are enriched with nutrients.

Temperature, pH, and total alkalinity were then excluded from the analyses to focus on salinity, K_d_, and nutrient concentrations, which are the parameters that varied the most between sites and periods. Dimension 1 of the Principal Component Analysis (PCA) was mainly driven by nutrient concentrations and salinity and explained more than 71 % of the variances (Fig. 2B). K_d_ allowed to differentiate sites and periods within dimension 2. For Kongsfjordneset, the weekly measured environmental parameters did not vary much. During week 25, at the beginning of the summer, all sites were grouped together with a similar high salinity. After week 25, nutrients increased and salinity decreased at all sites, except at Kongsfjordneset (arrow, Fig. 2). For Ossian Sars and Kongsbreen South sites, the K_d_ was particularly elevated on weeks 32 and 30 (early August), respectively. While K_d_ seemed not correlated with salinity and nutrient concentrations, these two high values could have driven the trend and the position of the arrow representing the K_d_.

### 2.2. eDNA

Our results show a high diversity of the faunal benthic community within the fjord with 44 taxa and Shannon indexes ranged between 2.17 to 3.18 (Fig. 1). A number of 17 macroalgae taxa was identified with Shannon indexes for macroalgae only going from 1.11 to 2.20. The list of taxa is provided in the supplementary material (Table S1). For both fauna and macroalgae, Shannon diversity indexes were significantly higher in Ossian Sars compared to the five other sites. This trend holds true when considering all periods were combined, as well as in June and July for fauna (Table S2). Ossian Sars showed both more taxa and a more balanced distribution between taxa than the other sites.

In June and July, the Kongsbreen South site, closer to the glacier, and French Bird Cliff, showed a significantly higher faunal and macroalgal diversity than Kongsfjordneset. Bloomstrand faunal diversity was also significatively higher than Kongsfjordneset both in June and July.

## 3. Discussion

In contrast to our initial hypotheses, we did not find a diversity gradient nor a correlation between the distribution of macroalgae and the underwater light regime nor a correlation of suspension feeders with the chl *a* or nutrient concentrations. Although we were not able to find a direct correlation between benthic diversity and environmental conditions, our results highlight some interesting observations. Apart from Kongsfjordneset, all other sites were affected by glacier melting. This was evidenced by an increase in nutrient levels observed from weeks 28 to 34 (early July to the end of August, Fig 2). In particular, during weeks 30 and 32 (early August), Ossian Sars and Kongsbreen South experienced a sharp increase in K_d_, which may coincide with the melting period of the glacier. The two sites the most influenced by glaciers also exhibited the highest levels of benthic diversity. In contrast, previous studies reported a decline in species richness from the outer to the inner part of Kongsfjorden (e.g. Kedra et al., 2010; Voronkov et al., 2013; Legeżyńska et al., 2017). Also, trends of decreasing diversity when getting closer to the glaciers have been observed in other glacial-impacted regions such as Hornsund fjord (Włodarska-Kowalczuk et al., 2013) and the Canadian Arctic (Farrow et al., 1983). Previous studies have suggested that the outer and central basins of fjords may be enriched with boreal species which may coexist with arctic and sub-arctic species already inhabiting these areas (e.g. Renaud et al., 2019). In Kongsfjorden, the absence of entrance sills allows shelf taxa to easily enter the fjord (Svendsen et al. 2002; Legezynska et al., 2017). Also, warming increases the discharge of freshwater enriched with inorganic particles into the fjord and the frequency of disturbance events such as ice scouring and sediment slides (Wlodarska-Kowalczuk et al. 2005). Although such increased disturbance may promote opportunistic species, it has been predicted that overall benthic diversity in these areas would decrease (Wlodarska-Kowalczuk et al. 2005; Al-Habahbeh et al. 2020). Hence, we would expect a diversity increase close to the entrance of the fjord, with more boreal species, and a decrease close to the glacier where disturbances events are more frequent. However, our results do not align with these expectations.

Kedra et al., (2010) have shown, using a sediment sampling method, that the diversity of the macrobenthic soft-bottom community in the middle and outer part of the Kongsfjorden decreased between 1997 and 2006. In contrast, the inner glacial bay, well-separated from the outer fjord by a chain of islands and a shallow sill (20 to 50 m; Svendsen et al. 2002) that acted as a barrier, remained as diverse, although it was still less than outer sites in this study. There was no major change in substrate type during this period, which therefore cannot explain this change. Kedra et al. (2010) put forward the hypothesis that the decline in diversity in the outer part of the fjord could be attributed to the decreasing influence of the Blomstrandbreen glacier which retreats quickly (Burton et al., 2016). With its retreat, its melting would no longer benefit the most distant sites. Indeed, glacier melting provides nutrients that support the growth of phytoplankton communities which form the basis of the food web (Piquet et al., 2014), and macroalgae, which are key habitat species in the ecosystem (Lebrun et al., 2022). Also, the presence of top predators in the Arctic coastal marine ecosystem is closely tied to the glacier front (Lydersen et al., 2014; Bouchard Marmen et al., 2017). In fact, top predators such as seabirds and sea lions migrate to the Arctic during the summer to live near glacier fronts, which underscores the importance of the nutritional richness these areas provide (Hop et al., 2002; Lydersen et al., 2014). Predators play a crucial role in regulating the ecosystem through top-down control, which may in turn further increase its diversity (Wessels et al., 2006; Bouchard Marmen et al., 2017). Hence, glaciers harbor key species and regulate the diversity of the benthic communities (Gutt et al., 2001).

The presence of top predators at the inner sites could explain our results. Seabirds are present in Ossian Sars, Kongsbreen South, Bloomstrand East, and French Bird Cliff (Varpe and Gabrielsen, 2022; pers. obs.). Sea Lions have been observed at Ossian Sars, Kongsbreen South, and Hansneset (Everett et al., 2018; pers. obs.). By identifying the presence of these top predators and their interactions with the benthic community, we established a trophic network diagram at each of the six sites (Fig. 3). Not surprisingly, the most diverse site also exhibited the most complex ecosystem with numerous trophic links and an homogenous repartition of taxa within and between the different trophic levels. Kongsbreen South, the site very close to the glacier, also presents a very complex ecosystem but with more heterogeneity between the first and the second levels. The presence of the bird colony at Ossian Sars could explain a better top-down regulation at that site than at Kongsbreen South. Conversely, Kongsfjordneset, the furthest site from the glaciers, only exhibits two heterogenous trophic levels. The presence of these top predators could therefore explain a greater diversity at Ossian Sars and Kongsbreen South, near the glaciers, in comparison with Kongsfjordneset, which no longer seems influenced by their melting.

**Figure 3:**
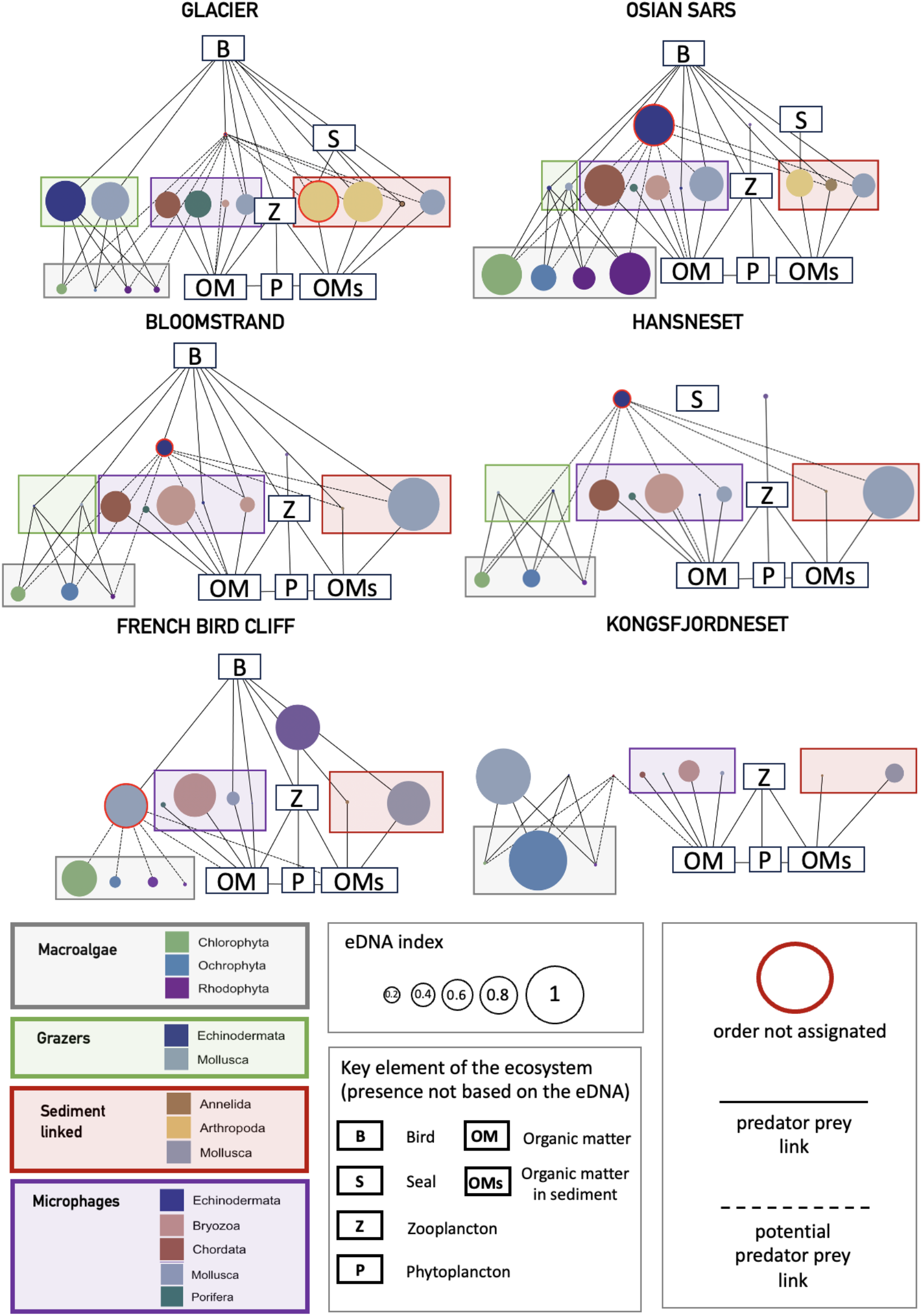
Trophic network diagram of each site based on the known predator-prey links between phyla, presence of predators, and the eDNA index.

The type of substrate (rocky vs. sediment) can also hold substantial importance in shaping the community composition (Lantuit et al., 2012; Filbee-Dexter et al., 2019). Hansneset and Kongsfjordneset feature rocky substrates, while the other sites predominantly consist of sediment accompanied with rocks. According to our findings, the rocky-substrate communities do not exhibit higher diversity. This align with the idea of predators being the key factor.

With warming, the retreat of glaciers could have a significant impact on the ecosystem. Specifically, the disappearance of the glacier front may lead to substantial changes in phytoplankton blooms and the properties of water masses in the fjord (Ardyna and Arrigo, 2020). These changes could, in turn, cause modifications in top predators’ presence and, hence, in ecosystem diversity. However, predicting these changes is very challenging.

It is important to note that like any other analysis method, eDNA sampling also has biases (Yates et al., 2019). During our exploration of Kongsfjordneset, the outermost site of the fjord, we observed a strong abundance of coralline algae while diving and using a drone, compared to other sites (Steeve Comeau pers. obs.). However, despite being present in the collected seawater samples from this site, the coralline algae exhibited a low eDNA index. Similarly, we observed a high presence of sea urchins while diving at Kongsfjordneset, but the eDNA index rather indicated a high abundance of mollusks, ochrophytes, and bryozoans. These differences in abundance determined by eDNA are difficult to interpret because although the eDNA index makes it possible to establish a value based on the number of sequences between 0 and 1, the number of sequences is not only linked to the abundance of a taxon. The eDNA index does not only reflect the abundance or biomass of a species or genus, but also the ability of its sequence to be amplified (Kelly et al., 2019). Additionally, the sequences can come from a dead organism or from the eggs or sperm, which can further bias the results. Therefore, while our results provide insights into the diversity of the studied sites, caution should be taken when interpreting them.

Furthermore, part of our replicates (40%) did not meet the quality standards required to be included in our study. These replicates were randomly of poor quality, i.e., not targeted on a specific date, site, or primer. This suggests that the quality issue was not due to a problem during sampling or extractions, which were done following the same protocols for all samples. Also, the PCR amplification was realized on all samples at the same time, with the same products. The poor per-base sequence quality obtained could have arisen from various sources, including errors during the sequencing process, which may lead to inaccuracies in base calling and subsequent data analysis. Also, DNA degradation before or during the sequencing procedure can significantly impact the quality of the obtained sequences, as damaged DNA molecules may result in incomplete or unreliable data. This limitation forced us to merge the remaining filtration and PCR replicates, thus reducing the robustness of our results.

The trend of decreasing diversity when approaching the interior of the fjord described previously could be reversed now as suggested by our results but further studies must validate it. Additional sampling of seawater for eDNA analysis throughout the fjord could determine the consistency of our results. Also, other biodiversity assessment methods like community sampling or photography of quadrats could validate our findings. Combined with stable isotope analysis to look into the trophic food web, the hypothesis of predator regulation could be tested. Overall, while our study provides some insight into the benthic diversity of Kongsfjorden, caution should be exercised when interpreting the results due to the limitations of our sampling method and the need for additional research.

## Acknowledgment

We are extremely grateful to Ryan Kelly for his contribution to experiment design and his invaluable assistance throughout the bioinformatics cleaning steps. Additionally, we express our sincere appreciation to Guri Gledis and Elizabeth Andruszkiewicz Allan for their help in the bioinformatic process. We thank Samir Alliouane and Cale Miller for their assistance during the data collection process. Cale Miller also contributed in helping to measure pH, alkalinity, and chl *a* concentrations. We thank the Molecular Biology Platform of Institut de la Mer de Villefranche (IMEV) that is supported by IR EMBRC-France, whose fundings are managed by the ANR within the Investments for Future Program under reference ANR-10-INBS-02. This study was supported by the EC2CO program of the Institut National des Sciences de l’Univers (INSU, CNRS). This study was conducted in the frame of the project FACE-IT (The Future of Arctic Coastal Ecosystems – Identifying Transitions in Fjord Systems and Adjacent Coastal Areas). FACE-IT has received funding from the European Union’s Horizon 2020 research and innovation programme under grant agreement No 869154.

## Author contributions

All authors designed the study and were involved in the fieldwork. AL, SC, FG, and JPG measured pH and alkalinity. AL performed the nutrient analysis. AL measured chl *a* concentrations. AL and SC performed the filtration of eDNA seawater samples. AL performed the extraction, PCR, and purification steps. AL performed the bioinformatics cleaning steps. AL analyzed the data and wrote the first draft of the manuscript which was accepted by all co-authors.

