## Supplementary material for "Benthic diversity along an Arctic fjord: which are the key factors?": Table S1

Table S1: eDNA-based records of phylum, order, and species across sites and sampling periods. The eDNA index was calculated using species-scale reads counts or, if not available (NA) the order or phylum scales.

| Month | Site | Phylum | Order | Species | nReads | eDNA index |
|---|---|---|---|---|---|---|
| June | Bloomstrand Est | Annelida | Sabellida | *Sabellidae sp. CMC02* | 4 | 1 |
| June | Bloomstrand Est | Annelida | Phyllodocida | NA | 7 | 0.551 |
| June | Bloomstrand Est | Bryozoa | Cyclostomatida | *Crisia eburnea* | 7 | 1 |
| June | Bloomstrand Est | Bryozoa | Cheilostomatida | NA | 33 | 1 |
| June | Bloomstrand Est | Chlorophyta | Ulotrichales | NA | 31 | 0.638 |
| June | Bloomstrand Est | Chordata | Stolidobranchia | NA | 5 | 1 |
| June | Bloomstrand Est | Chordata | Stolidobranchia | *Pelonaia corrugata* | 13 | 1 |
| June | Bloomstrand Est | Chordata | Phlebobranchia | NA | 2 | 0.571 |
| June | Bloomstrand Est | Echinodermata | Ophiuroidea sp. EAC01 | NA | 2 | 1 |
| June | Bloomstrand Est | Echinodermata | Echinoida | *Strongylocentrotus pallidus* | 3 | 0.034 |
| June | Bloomstrand Est | Mollusca | Littorinimorpha | *Lacuna vincta* | 23 | 1 |
| June | Bloomstrand Est | Mollusca | Adapedonta | *Hiatella arctica* | 61 | 0.103 |
| June | Bloomstrand Est | Mollusca | Myoida | *Hiatella sp. K HML-2015* | 34 | 0.446 |
| June | Bloomstrand Est | Ochrophyta | Ectocarpales | *Chordaria flagelliformis* | 3 | 1 |
| June | Bloomstrand Est | Ochrophyta | Ectocarpales | *Microspongium alariae* | 10 | 1 |
| June | Bloomstrand Est | Ochrophyta | Ectocarpales | NA | 16 | 1 |
| June | Bloomstrand Est | Ochrophyta | Ectocarpales | *Pylaiella littoralis* | 26 | 1 |
| June | Bloomstrand Est | Ochrophyta | Laminariales | NA | 198 | 0.296 |
| June | Bloomstrand Est | Porifera | Suberitida | NA | 9 | 1 |
| June | Bloomstrand Est | Porifera | Bubarida | NA | 235 | 0.486 |
| June | French Bird Cliff | Annelida | Capitellida | *Arenicola marina* | 5 | 1 |
| June | French Bird Cliff | Annelida | Phyllodocida | NA | 23 | 1 |
| June | French Bird Cliff | Arthropoda | Decapoda | *Hyas araneus* | 119 | 1 |
| June | French Bird Cliff | Bryozoa | Ctenostomatida | *Vesicularia spinosa* | 2 | 1 |
| June | French Bird Cliff | Chlorophyta | Ulotrichales | NA | 88 | 1 |
| June | French Bird Cliff | Cnidaria | Siphonophorae | *Apolemia sp. BO-2009* | 143 | 1 |
| June | French Bird Cliff | Cnidaria | Leptothecata | *Calycella syringa* | 3 | 1 |
| June | French Bird Cliff | Cnidaria | Anthoathecata | *Sarsia princeps* | 45 | 1 |
| June | French Bird Cliff | Cnidaria | Anthoathecata | *Eudendrium racemosum* | 2 | 0.075 |
| June | French Bird Cliff | Echinodermata | Amphilepidida | NA | 13 | 1 |
| June | French Bird Cliff | Echinodermata | Ophiurida | *Ophiopholis aculeata* | 59 | 0.097 |
| June | French Bird Cliff | Mollusca | Hygrophila | *Biomphalaria oligoza* | 101 | 1 |
| June | French Bird Cliff | Mollusca | Veneroida | *Macoma calcarea* | 51 | 1 |
| June | French Bird Cliff | Mollusca | Cardiida | NA | 9 | 1 |
| June | French Bird Cliff | Mollusca | Myida | NA | 21 | 1 |
| June | French Bird Cliff | Mollusca | Adapedonta | *Hiatella arctica* | 96 | 0.089 |
| June | French Bird Cliff | Mollusca | Myoida | *Hiatella sp. K HML-2015* | 47 | 0.340 |
| June | French Bird Cliff | Ochrophyta | Ectocarpales | *cinetosporaceae sp. 2 AP-2014* | 7 | 1 |
| June | French Bird Cliff | Ochrophyta | Laminariales | *Chorda filum* | 7 | 1 |
| June | French Bird Cliff | Ochrophyta | Ectocarpales | *Cladosiphon zosterae* | 25 | 1 |
| June | French Bird Cliff | Ochrophyta | Laminariales | NA | 19 | 0.015 |
| June | French Bird Cliff | Ochrophyta | Ectocarpales | *Chordaria flagelliformis* | 4 | 0.735 |
| June | French Bird Cliff | Porifera | Suberitida | *Halichondria panicea* | 22 | 0.300 |
| June | French Bird Cliff | Porifera | Bubarida | NA | 364 | 0.415 |
| June | French Bird Cliff | Rhodophyta | Palmariales | *Devaleraea ramentacea* | 4 | 1 |
| June | French Bird Cliff | Rhodophyta | Bangiales | *Pyropia haitanensis* | 6 | 1 |
| June | French Bird Cliff | Rhodophyta | Palmariales | NA | 8 | 0.227 |
| June | French Bird Cliff | Rhodophyta | NA | NA | 15 | 0.253 |
| June | Glacier | Annelida | Spionida | *Cirratulus cirratus* | 31 | 1 |
| June | Glacier | Annelida | Phyllodocida | *Glycera capitata* | 2 | 1 |
| June | Glacier | Annelida | Spionida | *Polydora onagawaensis* | 2 | 0.000 |
| June | Glacier | Annelida | Terebellida | *Amphitrite cirrata* | 9 | 0.074 |
| June | Glacier | Arthropoda | Cyclopoida | *cilostomatoida sp. 09PROBE-08* | 10 | 1 |
| June | Glacier | Bryozoa | Cheilostomatida | *Cauloramphus magnus* | 4 | 0.974 |
| June | Glacier | Chordata | Stolidobranchia | *Halocynthia pyriformis* | 2 | 0.028 |
| June | Glacier | Cnidaria | Siphonophorae | *Apolemia sp. BO-2009* | 18 | 0.034 |
| June | Glacier | Echinodermata | Camarodonta | NA | 2 | 0.053 |
| June | Glacier | Echinodermata | Echinoida | *Strongylocentrotus pallidus* | 264 | 0.454 |
| June | Glacier | Mollusca | Lepidopleurida | NA | 7 | 1 |
| June | Glacier | Mollusca | Nudibranchia | *Zelentia willowsi* | 4 | 1 |
| June | Glacier | Mollusca | Hygrophila | *Biomphalaria sp. GC-2010* | 3 | 0.012 |
| June | Glacier | Mollusca | Myoida | *Hiatella sp. K HML-2015* | 94 | 0.185 |
| June | Glacier | Mollusca | Chitonida | *Tonicella undocaerulea* | 698 | 0.377 |
| June | Glacier | Mollusca | Littorinimorpha | *Lacuna vincta* | 68 | 0.445 |
| June | Glacier | Mollusca | Adapedonta | *Hiatella arctica* | 2916 | 0.743 |
| June | Glacier | Ochrophyta | Ectocarpales | *Chordariaceae sp. 4nov* | 6 | 1 |
| June | Glacier | Ochrophyta | Ectocarpales | *Pylaiella washingtoniensis* | 6 | 1 |
| June | Glacier | Ochrophyta | Laminariales | NA | 25 | 0.005 |
| June | Glacier | Ochrophyta | Ectocarpales | *Microspongium alariae* | 6 | 0.090 |
| June | Glacier | Ochrophyta | Ectocarpales | *Chordaria flagelliformis* | 6 | 0.301 |
| June | Glacier | Porifera | Suberitida | *Halichondria panicea* | 27 | 0.100 |
| June | Glacier | Porifera | Bubarida | NA | 502 | 0.156 |
| June | Glacier | Rhodophyta | NA | NA | 16 | 1 |
| June | Glacier | Rhodophyta | Palmariales | NA | 5 | 0.043 |
| June | Glacier | Rhodophyta | Palmariales | *Rhodophysema kjellmanii* | 4 | 0.123 |
| June | Glacier | Rhodophyta | Palmariales | *Devaleraea ramentacea* | 8 | 0.290 |
| June | Hansneset | Annelida | olychaeta sp. 09PROBE-0831 | NA | 6 | 1 |
| June | Hansneset | Annelida | Spionida | *Polydora onagawaensis* | 264 | 1 |
| June | Hansneset | Mollusca | Veneroida | *Tridacna squamosa* | 2 | 1 |
| June | Hansneset | Mollusca | Myoida | *Hiatella sp. K HML-2015* | 4 | 0.090 |
| June | Hansneset | Mollusca | Littorinimorpha | *Lacuna vincta* | 3 | 0.224 |
| June | Hansneset | Ochrophyta | Ectocarpales | *Dictyosiphon foeniculaceus* | 2 | 1 |
| June | Hansneset | Porifera | Bubarida | NA | 138 | 0.492 |
| June | Kongsfjordneset | Annelida | Phyllodocida | NA | 13 | 1 |
| June | Kongsfjordneset | Annelida | NA | *Arenicola marina* | 3 | 0.041 |
| June | Kongsfjordneset | Annelida | NA | NA | 2 | 0.366 |
| June | Kongsfjordneset | Bryozoa | Cyclostomatida | *Crisia eburnea* | 7 | 0.357 |
| June | Kongsfjordneset | Chlorophyta | Ulvales | *Ulvella leptochaete* | 4 | 1 |
| June | Kongsfjordneset | Chlorophyta | Ulotrichales | NA | 26 | 0.191 |
| June | Kongsfjordneset | Chordata | Stolidobranchia | *Dendrodoa grossularia* | 2 | 0.195 |
| June | Kongsfjordneset | Mollusca | Chitonida | *Tonicella undocaerulea* | 12 | 0.015 |
| June | Kongsfjordneset | Mollusca | Adapedonta | *Hiatella arctica* | 965 | 0.584 |
| June | Kongsfjordneset | Ochrophyta | Desmarestiales | NA | 19 | 1 |
| June | Kongsfjordneset | Ochrophyta | Laminariales | NA | 925 | 0.495 |
| June | Kongsfjordneset | Ochrophyta | Laminariales | *Halosiphon tomentosus* | 35 | 0.631 |
| June | Kongsfjordneset | Porifera | Dendroceratida | *Halisarca dujardinii* | 6 | 1 |
| June | Ossian Sars | Annelida | NA | *Arenicola marina* | 4 | 1 |
| June | Ossian Sars | Annelida | Spionida | *Chaetozone setosa* | 35 | 1 |
| June | Ossian Sars | Annelida | NA | NA | 94 | 1 |
| June | Ossian Sars | Annelida | Sabellida | *Simplaria potswaldi* | 4 | 1 |
| June | Ossian Sars | Annelida | Terebellida | *Spinosphaera hutchingsae* | 4 | 1 |
| June | Ossian Sars | Annelida | Sabellida | NA | 33 | 0.245 |
| June | Ossian Sars | Annelida | Phyllodocida | NA | 17 | 0.682 |
| June | Ossian Sars | Bryozoa | Cyclostomatida | *Annectocyma tubulosa* | 2 | 1 |
| June | Ossian Sars | Bryozoa | Cheilostomatida | NA | 5 | 0.027 |
| June | Ossian Sars | Bryozoa | Cyclostomatida | *Crisia eburnea* | 2 | 0.052 |
| June | Ossian Sars | Chlorophyta | Ulotrichales | NA | 124 | 0.467 |
| June | Ossian Sars | Chordata | Stolidobranchia | *Dendrodoa grossularia* | 20 | 1 |
| June | Ossian Sars | Chordata | Stolidobranchia | NA | 7 | 0.140 |
| June | Ossian Sars | Chordata | Stolidobranchia | *Pelonaia corrugata* | 10 | 0.256 |
| June | Ossian Sars | Cnidaria | Leptothecata | *Obelia longissima* | 5 | 1 |
| June | Ossian Sars | Echinodermata | Camarodonta | NA | 31 | 1 |
| June | Ossian Sars | Echinodermata | Amphilepidida | NA | 2 | 0.051 |
| June | Ossian Sars | Mollusca | Trochida | *Margarites helicinus* | 11 | 1 |
| June | Ossian Sars | Mollusca | Littorinimorpha | NA | 39 | 1 |
| June | Ossian Sars | Mollusca | Myoida | *Hiatella sp. K HML-2015* | 9 | 0.021 |
| June | Ossian Sars | Mollusca | Chitonida | *Tonicella undocaerulea* | 39 | 0.025 |
| June | Ossian Sars | Mollusca | Adapedonta | *Hiatella arctica* | 644 | 0.199 |
| June | Ossian Sars | Mollusca | Myida | NA | 17 | 0.268 |
| June | Ossian Sars | Mollusca | Cardiida | NA | 26 | 0.958 |
| June | Ossian Sars | Ochrophyta | Ectocarpales | NA | 18 | 0.205 |
| June | Ossian Sars | Ochrophyta | Desmarestiales | NA | 20 | 0.538 |

| Month | Site | Phylum | Order | Species | nReads | eDNA index |
|---|---|---|---|---|---|---|
| July | Ossian Sars | Porifera | Leucosolenida | *Sycon abyssale* | 11 | 1 |
| July | Ossian Sars | Porifera | Suberitida | *Halichondria panicea* | 10 | 0.033 |
| July | Ossian Sars | Porifera | Bubarida | NA | 640 | 0.178 |
| July | Ossian Sars | Rhodophyta | Palmariales | *Devaleraea ramentacea* | 72 | 1 |
| July | Ossian Sars | Rhodophyta | Palmariales | NA | 129 | 1 |
| July | Ossian Sars | Rhodophyta | Ceramiales | *Savoiea arctica* | 3 | 0.067 |
| July | Ossian Sars | Rhodophyta | Hapalidiales | NA | 21 | 0.737 |
| August | Bloomstrand Est | Annelida | NA | NA | 12 | 0.074 |
| August | Bloomstrand Est | Chordata | Stolidobranchia | NA | 3 | 0.064 |
| August | Bloomstrand Est | Cnidaria | Siphonophorae | *Apolemia sp. BO-2009* | 44 | 0.059 |
| August | Bloomstrand Est | Cnidaria | Anthoathecata | *Eudendrium racemosum* | 18 | 0.132 |
| August | Bloomstrand Est | Echinodermata | Echinoida | *Strongylocentrotus pallidus* | 6 | 0.007 |
| August | Bloomstrand Est | Echinodermata | Ophiurida | *Ophiopholis aculeata* | 31 | 0.009 |
| August | Bloomstrand Est | Mollusca | Chitonida | *Tonicella undocaerulea* | 54 | 0.020 |
| August | Bloomstrand Est | Mollusca | Veneroida | *Macoma calcarea* | 8 | 0.030 |
| August | Bloomstrand Est | Mollusca | Myoida | *Hiatella sp. K HML-2015* | 52 | 0.073 |
| August | Bloomstrand Est | Mollusca | Hygrophila | *Biomphalaria oligoza* | 46 | 0.088 |
| August | Bloomstrand Est | Mollusca | Myida | NA | 65 | 0.601 |
| August | Bloomstrand Est | Mollusca | Adapedonta | *Hiatella arctica* | 5481 | 0.995 |
| August | Bloomstrand Est | Ochrophyta | Laminariales | *Halosiphon tomentosus* | 185 | 1 |
| August | Bloomstrand Est | Ochrophyta | Ectocarpales | *Cladosiphon zosterae* | 4 | 0.031 |
| August | Bloomstrand Est | Ochrophyta | Laminariales | NA | 223 | 0.035 |
| August | Bloomstrand Est | Ochrophyta | Ectocarpales | NA | 29 | 0.115 |
| August | Bloomstrand Est | Ochrophyta | Ectocarpales | *Pylaiella littoralis* | 28 | 0.194 |
| August | Bloomstrand Est | Porifera | Bubarida | NA | 447 | 0.099 |
| August | French Bird Cliff | Annelida | NA | NA | 58 | 0.280 |
| August | French Bird Cliff | Annelida | Phyllodocida | NA | 42 | 0.285 |
| August | French Bird Cliff | Bryozoa | Cheilostomatida | *Eucratea loricata* | 64 | 1 |
| August | French Bird Cliff | Chlorophyta | NA | NA | 114 | 0.024 |
| August | French Bird Cliff | Chlorophyta | Ulotrichales | NA | 14 | 0.505 |
| August | French Bird Cliff | Cnidaria | Anthoathecata | *Eudendrium racemosum* | 172 | 1 |
| August | French Bird Cliff | Mollusca | Adapedonta | *Hiatella arctica* | 6745 | 0.967 |
| August | French Bird Cliff | Ochrophyta | Ectocarpales | *Pylaiella littoralis* | 95 | 0.084 |
| August | French Bird Cliff | Ochrophyta | Laminariales | NA | 1173 | 0.148 |
| August | French Bird Cliff | Ochrophyta | Desmarestiales | NA | 14 | 0.174 |
| August | French Bird Cliff | Ochrophyta | Ectocarpales | NA | 16 | 0.309 |
| August | French Bird Cliff | Rhodophyta | Palmariales | NA | 23 | 0.111 |
| August | Glacier | Annelida | NA | NA | 3 | 0.095 |
| August | Glacier | Arthropoda | Euphausiacea | NA | 211 | 1 |
| August | Glacier | Chlorophyta | Ulotrichales | NA | 30 | 0.339 |
| August | Glacier | Cnidaria | Siphonophorae | *Apolemia sp. BO-2009* | 107 | 0.745 |
| August | Glacier | Mollusca | Hygrophila | *Biomphalaria sp. GC-2010* | 67 | 1 |
| August | Glacier | Mollusca | Adapedonta | *Hiatella arctica* | 37 | 0.034 |
| August | Glacier | Mollusca | Chitonida | *Tonicella undocaerulea* | 33 | 0.065 |
| August | Glacier | Mollusca | Myoida | *Hiatella sp. K HML-2015* | 16 | 0.115 |
| August | Glacier | Ochrophyta | Laminariales | NA | 48 | 0.039 |
| August | Glacier | Porifera | Bubarida | NA | 750 | 0.854 |
| August | Glacier | Rhodophyta | Ceramiales | *Rhodomela confervoides* | 2 | 1 |
| August | Glacier | Rhodophyta | Palmariales | NA | 8 | 0.252 |
| August | Hansneset | Annelida | Phyllodocida | *Pholoe baltica* | 8 | 1 |
| August | Hansneset | Annelida | Spionida | *Polydora onagawaensis* | 21 | 0.009 |
| August | Hansneset | Annelida | Terebellida | *Amphitrite cirrata* | 33 | 0.372 |
| August | Hansneset | Annelida | Sabellida | *Sabellidae sp. CMC02* | 17 | 0.875 |
| August | Hansneset | Arthropoda | Decapoda | *Hyas araneus* | 14 | 0.043 |
| August | Hansneset | Bryozoa | Cheilostomatida | *Cauloramphus magnus* | 3 | 1 |
| August | Hansneset | Bryozoa | Cheilostomatida | *Dendrobeania murrayana* | 3 | 1 |
| August | Hansneset | Bryozoa | Cheilostomatida | *Tricellaria ternata* | 7 | 1 |
| August | Hansneset | Chlorophyta | Ulotrichales | NA | 21 | 0.089 |
| August | Hansneset | Chordata | Stolidobranchia | *Halocynthia pyriformis* | 18 | 0.353 |
| August | Hansneset | Cnidaria | Siphonophorae | *Apolemia sp. BO-2009* | 264 | 0.688 |
| August | Hansneset | Echinodermata | Ophiurida | *Ophiopholis aculeata* | 1618 | 1 |
| August | Hansneset | Echinodermata | Echinoida | *Strongylocentrotus pallidus* | 183 | 0.430 |

| Month | Site | Phylum | Order | Species | nReads | eDNA index |
|---|---|---|---|---|---|---|
| August | Hansneset | Echinodermata | Amphilepidida | NA | 24 | 0.688 |
| August | Hansneset | Mollusca | Adapedonta | *Hiatella arctica* | 110 | 0.038 |
| August | Hansneset | Mollusca | Myoida | *Hiatella sp. K HML-2015* | 21 | 0.056 |
| August | Hansneset | Ochrophyta | Ectocarpales | *Chordaria chordaeformis* | 2 | 1 |
| August | Hansneset | Ochrophyta | Laminariales | NA | 311 | 0.095 |
| August | Hansneset | Ochrophyta | Ectocarpales | *Microspongium alariae* | 17 | 0.350 |
| August | Hansneset | Porifera | Poecilosclerida | *Crella elegans* | 12 | 1 |
| August | Hansneset | Porifera | Suberitida | *Halichondria panicea* | 196 | 1 |
| August | Hansneset | Porifera | Bubarida | NA | 535 | 0.228 |
| August | Hansneset | Rhodophyta | Acrochaetiales | *Acrochaetium secundatum* | 2 | 1 |
| August | Hansneset | Rhodophyta | Ceramiales | *Savoiea arctica* | 29 | 1 |
| August | Hansneset | Rhodophyta | Palmariales | NA | 19 | 0.224 |
| August | Hansneset | Rhodophyta | Palmariales | *Devaleraea ramentacea* | 17 | 0.360 |
| August | Kongsfjordneset | Annelida | NA | *Arenicola marina* | 3 | 1 |
| August | Kongsfjordneset | Annelida | Sabellida | *Chitinopoma serrula* | 3 | 0.054 |
| August | Kongsfjordneset | Annelida | Spionida | *Cirratulus cirratus* | 40 | 0.107 |
| August | Kongsfjordneset | Annelida | Phyllodocida | NA | 19 | 0.200 |
| August | Kongsfjordneset | Annelida | NA | NA | 13 | 0.613 |
| August | Kongsfjordneset | Bryozoa | Ctenostomatida | *Amathia gracilis* | 61 | 1 |
| August | Kongsfjordneset | Bryozoa | Cheilostomatida | *Eucratea loricata* | 3 | 0.039 |
| August | Kongsfjordneset | Chlorophyta | NA | NA | 2 | 0.007 |
| August | Kongsfjordneset | Chordata | Stolidobranchia | NA | 4 | 0.057 |
| August | Kongsfjordneset | Chordata | Stolidobranchia | *Halocynthia pyriformis* | 10 | 0.068 |
| August | Kongsfjordneset | Cnidaria | Anthoathecata | *Sarsia princeps* | 4 | 0.011 |
| August | Kongsfjordneset | Echinodermata | Echinoida | *Strongylocentrotus pallidus* | 3 | 0.002 |
| August | Kongsfjordneset | Echinodermata | Camarodonta | NA | 7 | 0.088 |
| August | Kongsfjordneset | Mollusca | Chitonida | *Tonicella undocaerulea* | 3882 | 1 |
| August | Kongsfjordneset | Mollusca | Littorinimorpha | *Lacuna vincta* | 5 | 0.015 |
| August | Kongsfjordneset | Mollusca | Myoida | *Hiatella sp. K HML-2015* | 71 | 0.066 |
| August | Kongsfjordneset | Mollusca | Adapedonta | *Hiatella arctica* | 5485 | 0.665 |
| August | Kongsfjordneset | Ochrophyta | Laminariales | NA | 119 | 0.012 |
| August | Kongsfjordneset | Ochrophyta | Desmarestiales | NA | 71 | 0.748 |
| August | Kongsfjordneset | Porifera | Bubarida | NA | 263 | 0.038 |
| August | Kongsfjordneset | Porifera | Haplosclerida | NA | 4 | 0.521 |
| August | Kongsfjordneset | Rhodophyta | Hapalidiales | *Leptophytum laeve* | 7 | 1 |
| August | Kongsfjordneset | Rhodophyta | Palmariales | NA | 3 | 0.012 |
| August | Ossian Sars | Annelida | Spionida | *Polydora onagawaensis* | 16 | 1 |
| August | Ossian Sars | Annelida | NA | *Arenicola marina* | 4 | 0.005 |
| August | Ossian Sars | Annelida | Sabellida | *Paradexiospira vitrea* | 5 | 0.275 |
| August | Ossian Sars | Annelida | Phyllodocida | NA | 23 | 0.566 |
| August | Ossian Sars | Bryozoa | Cyclostomatida | *Crisia eburnea* | 12 | 0.260 |
| August | Ossian Sars | Bryozoa | Cheilostomatida | *Tricellaria circumternata* | 14 | 0.787 |
| August | Ossian Sars | Chlorophyta | Ulotrichales | NA | 61 | 0.190 |
| August | Ossian Sars | Chordata | Stolidobranchia | *Pelonaia corrugata* | 13 | 0.151 |
| August | Ossian Sars | Chordata | Stolidobranchia | *Dendrodoa grossularia* | 4 | 0.165 |
| August | Ossian Sars | Cnidaria | Siphonophorae | *Apolemia sp. BO-2009* | 11 | 0.021 |
| August | Ossian Sars | Echinodermata | Apodida | NA | 89 | 1 |
| August | Ossian Sars | Mollusca | Adapedonta | *Hiatella arctica* | 3886 | 1 |
| August | Ossian Sars | Mollusca | Chitonida | *Tonicella undocaerulea* | 35 | 0.019 |
| August | Ossian Sars | Mollusca | Cardiida | NA | 32 | 0.978 |
| August | Ossian Sars | Ochrophyta | Ectocarpales | NA | 11 | 0.052 |
| August | Ossian Sars | Ochrophyta | Laminariales | NA | 414 | 0.094 |
| August | Ossian Sars | Ochrophyta | Ectocarpales | *Pylaiella littoralis* | 9 | 0.104 |
| August | Ossian Sars | Ochrophyta | Desmarestiales | NA | 18 | 0.402 |
| August | Ossian Sars | Porifera | Bubarida | NA | 61 | 0.019 |
| August | Ossian Sars | Porifera | Suberitida | NA | 5 | 0.084 |
| August | Ossian Sars | Rhodophyta | Hapalidiales | NA | 2 | 0.079 |
| August | Ossian Sars | Rhodophyta | Palmariales | NA | 29 | 0.252 |

| Month | Site | Phylum | Order | Species | nReads | eDNA index |
|---|---|---|---|---|---|---|
| June | Ossian Sars | Ochrophyta | Laminariales | NA | 2295 | 0.628 |
| June | Ossian Sars | Ochrophyta | Laminariales | *Halosiphon tomentosus* | 95 | 0.877 |
| June | Ossian Sars | Porifera | Haplosclerida | NA | 3 | 1 |
| June | Ossian Sars | Porifera | Suberitida | *Halichondria panicea* | 2 | 0.009 |
| June | Ossian Sars | Porifera | Bubarida | NA | 187 | 0.070 |
| June | Ossian Sars | Porifera | Suberitida | NA | 14 | 0.284 |
| June | Ossian Sars | Rhodophyta | Hapalidiales | NA | 21 | 1 |
| June | Ossian Sars | Rhodophyta | Palmariales | NA | 90 | 0.243 |
| June | Ossian Sars | Rhodophyta | Ceramiales | *Rhodomela confervoides* | 4 | 0.665 |
| June | Ossian Sars | Rhodophyta | NA | NA | 11 | 0.945 |
| July | Bloomstrand Est | Annelida | Phyllodocida | NA | 49 | 0.410 |
| July | Bloomstrand Est | Bryozoa | Cheilostomatida | *Tricellaria circumternata* | 5 | 0.196 |
| July | Bloomstrand Est | Bryozoa | Cyclostomatida | *Crisia eburnea* | 26 | 0.395 |
| July | Bloomstrand Est | Chlorophyta | Ulotrichales | NA | 45 | 0.098 |
| July | Bloomstrand Est | Chordata | Stolidobranchia | *Dendrodoa grossularia* | 5 | 0.145 |
| July | Bloomstrand Est | Chordata | Stolidobranchia | *Pelonaia corrugata* | 18 | 0.147 |
| July | Bloomstrand Est | Echinodermata | Camarodonta | NA | 6 | 0.112 |
| July | Bloomstrand Est | Echinodermata | Apodida | NA | 29 | 0.228 |
| July | Bloomstrand Est | Mollusca | Chitonida | *Tonicella undocaerulea* | 6 | 0.002 |
| July | Bloomstrand Est | Mollusca | Adapedonta | *Hiatella arctica* | 4389 | 0.791 |
| July | Bloomstrand Est | Ochrophyta | Ectocarpales | NA | 18 | 0.057 |
| July | Bloomstrand Est | Ochrophyta | Laminariales | *Halosiphon tomentosus* | 22 | 0.118 |
| July | Bloomstrand Est | Ochrophyta | Ectocarpales | *Pylaiella littoralis* | 14 | 0.119 |
| July | Bloomstrand Est | Ochrophyta | Desmarestiales | NA | 13 | 0.203 |
| July | Bloomstrand Est | Ochrophyta | Laminariales | NA | 2066 | 0.328 |
| July | Bloomstrand Est | Porifera | Bubarida | NA | 58 | 0.012 |
| July | Bloomstrand Est | Porifera | Suberitida | NA | 10 | 0.118 |
| July | Bloomstrand Est | Rhodophyta | Hapalidiales | NA | 9 | 0.249 |
| July | French Bird Cliff | Annelida | Phyllodocida | NA | 4 | 0.043 |
| July | French Bird Cliff | Annelida | NA | NA | 2 | 0.061 |
| July | French Bird Cliff | Annelida | Terebellida | *Amphitrite cirrata* | 3 | 0.118 |
| July | French Bird Cliff | Bryozoa | Cyclostomatida | *Crisia eburnea* | 6 | 0.321 |
| July | French Bird Cliff | Bryozoa | Cheilostomatida | *Eucratea loricata* | 5 | 0.345 |
| July | French Bird Cliff | Chlorophyta | Ulotrichales | NA | 82 | 0.058 |
| July | French Bird Cliff | Chlorophyta | NA | NA | 3 | 0.632 |
| July | French Bird Cliff | Chordata | Stolidobranchia | *Pelonaia corrugata* | 8 | 0.230 |
| July | French Bird Cliff | Cnidaria | Stauromedusae | *Lucernaria quadricornis* | 2 | 1 |
| July | French Bird Cliff | Echinodermata | Amphilepidida | NA | 19 | 0.992 |
| July | French Bird Cliff | Mollusca | Chitonida | *Tonicella undocaerulea* | 12 | 0.016 |
| July | French Bird Cliff | Mollusca | Myida | NA | 7 | 0.226 |
| July | French Bird Cliff | Mollusca | Adapedonta | *Hiatella arctica* | 1179 | 0.748 |
| July | French Bird Cliff | Ochrophyta | Ectocarpales | NA | 12 | 0.100 |
| July | French Bird Cliff | Ochrophyta | Laminariales | NA | 383 | 0.214 |
| July | French Bird Cliff | Ochrophyta | Ectocarpales | *Pylaiella littoralis* | 7 | 0.281 |
| July | French Bird Cliff | Ochrophyta | Laminariales | *Halosiphon tomentosus* | 22 | 0.415 |
| July | French Bird Cliff | Porifera | Poecilosclerida | *Myxilla incrustans* | 9 | 1 |
| July | French Bird Cliff | Porifera | Bubarida | NA | 130 | 0.100 |
| July | French Bird Cliff | Porifera | Suberitida | NA | 3 | 0.124 |
| July | French Bird Cliff | Porifera | Dendroceratida | *Halisarca dujardinii* | 3 | 0.523 |
| July | French Bird Cliff | Rhodophyta | Palmariales | NA | 26 | 0.559 |
| July | Glacier | Annelida | Spionida | *Polydora onagawaensis* | 16 | 0.030 |
| July | Glacier | Annelida | Phyllodocida | NA | 2 | 0.137 |
| July | Glacier | Annelida | Terebellida | *Amphitrite cirrata* | 9 | 0.432 |
| July | Glacier | Annelida | Capitellida | *Arenicola marina* | 2 | 0.634 |
| July | Glacier | Arthropoda | Balanomorpha | *Chthamalus fragilis* | 2 | 1 |
| July | Glacier | Bryozoa | Cyclostomatida | *Crisia eburnea* | 4 | 0.500 |
| July | Glacier | Chlorophyta | Ulvales | NA | 4 | 1 |
| July | Glacier | Chordata | Stolidobranchia | *Halocynthia pyriformis* | 12 | 1 |
| July | Glacier | Chordata | Phlebobranchia | NA | 4 | 1 |
| July | Glacier | Chordata | Stolidobranchia | *Pelonaia corrugata* | 6 | 0.403 |
| July | Glacier | Echinodermata | Dendrochirotida | *Cucumaria frondosa* | 3 | 1 |
| July | Glacier | Echinodermata | Echinoida | *Strongylocentrotus pallidus* | 100 | 1 |

| Month | Site | Phylum | Order | Species | nReads | eDNA index |
|---|---|---|---|---|---|---|
| July | Glacier | Mollusca | Hygrophila | *Biomphalaria oligoza* | 6 | 0.094 |
| July | Glacier | Mollusca | Veneroida | *Macoma calcarea* | 4 | 0.124 |
| July | Glacier | Mollusca | Myoida | *Hiatella sp. K HML-2015* | 20 | 0.229 |
| July | Glacier | Mollusca | Littorinimorpha | *Lacuna vincta* | 9 | 0.342 |
| July | Glacier | Ochrophyta | Ectocarpales | *Acinetosporaceae sp. 1 AP-2016* | 2 | 1 |
| July | Glacier | Ochrophyta | Laminariales | NA | 122 | 0.159 |
| July | Glacier | Ochrophyta | Ectocarpales | *Pylaiella littoralis* | 17 | 0.572 |
| July | Glacier | Porifera | Suberitida | *Halichondria panicea* | 18 | 0.390 |
| July | Glacier | Porifera | Suberitida | NA | 8 | 0.777 |
| July | Glacier | Porifera | Bubarida | NA | 450 | 0.815 |
| July | Glacier | Rhodophyta | Corallinales | NA | 5 | 1 |
| July | Hansneset | Annelida | Spionida | *Polydora onagawaensis* | 99 | 0.118 |
| July | Hansneset | Chlorophyta | Ulotrichales | NA | 6 | 0.067 |
| July | Hansneset | Cnidaria | Siphonophorae | *Apolemia sp. BO-2009* | 60 | 0.413 |
| July | Hansneset | Echinodermata | Ophiurida | *Ophiopholis aculeata* | 7 | 0.011 |
| July | Hansneset | Echinodermata | Echinoida | *Strongylocentrotus pallidus* | 19 | 0.118 |
| July | Hansneset | Mollusca | Myoida | *Hiatella sp. K HML-2015* | 140 | 1 |
| July | Hansneset | Mollusca | Adapedonta | *Hiatella arctica* | 30 | 0.027 |
| July | Hansneset | Mollusca | Hygrophila | *Biomphalaria oligoza* | 51 | 0.497 |
| July | Hansneset | Mollusca | Myida | NA | 11 | 0.516 |
| July | Hansneset | Ochrophyta | Ectocarpales | *Microspongium alariae* | 12 | 0.652 |
| July | Hansneset | Porifera | Bubarida | NA | 888 | 1 |
| July | Hansneset | Porifera | Suberitida | *Halichondria panicea* | 4 | 0.053 |
| July | Kongsfjordneset | Annelida | NA | *Scalibregma inflatum* | 7 | 1 |
| July | Kongsfjordneset | Annelida | NA | NA | 209 | 0.000 |
| July | Kongsfjordneset | Annelida | Spionida | *Polydora onagawaensis* | 6 | 0.478 |
| July | Kongsfjordneset | Arthropoda | NA | NA | 7 | 1 |
| July | Kongsfjordneset | Chlorophyta | NA | NA | 484 | 1 |
| July | Kongsfjordneset | Mollusca | Adapedonta | *Hiatella arctica* | 499 | 0.033 |
| July | Kongsfjordneset | Ochrophyta | Laminariales | NA | 16949 | 1 |
| July | Kongsfjordneset | Ochrophyta | Ectocarpales | NA | 23 | 0.056 |
| July | Kongsfjordneset | Porifera | Bubarida | NA | 129 | 0.010 |
| July | Ossian Sars | Annelida | NA | *Arenicola marina* | 5 | 1 |
| July | Ossian Sars | Annelida | Terebellida | *Spinosphaera hutchingsae* | 4 | 0.379 |
| July | Ossian Sars | Annelida | Sabellida | NA | 17 | 0.549 |
| July | Ossian Sars | Annelida | Phyllodocida | NA | 92 | 0.629 |
| July | Ossian Sars | Annelida | NA | NA | 70 | 0.737 |
| July | Ossian Sars | Annelida | Terebellida | *Amphitrite cirrata* | 135 | 0.978 |
| July | Ossian Sars | Arthropoda | Decapoda | *Hyas araneus* | 3 | 0.006 |
| July | Ossian Sars | Arthropoda | Euphausiacea | NA | 281 | 0.326 |
| July | Ossian Sars | Bryozoa | Cheilostomatida | *Tricellaria circumternata* | 20 | 1 |
| July | Ossian Sars | Chlorophyta | Ulvales | *Ulvella leptochaete* | 4 | 0.154 |
| July | Ossian Sars | Chlorophyta | Ulotrichales | NA | 77 | 0.214 |
| July | Ossian Sars | Chlorophyta | Ulvales | NA | 4 | 0.377 |
| July | Ossian Sars | Chordata | Stolidobranchia | *Dendrodoa grossularia* | 23 | 0.848 |
| July | Ossian Sars | Cnidaria | Leptothecata | *Orthopyxis integra* | 2 | 1 |
| July | Ossian Sars | Cnidaria | Anthoathecata | *Zanclea implexa* | 2 | 1 |
| July | Ossian Sars | Cnidaria | Anthoathecata | *Sarsia princeps* | 3 | 0.016 |
| July | Ossian Sars | Cnidaria | Siphonophorae | *Apolemia sp. BO-2009* | 24 | 0.041 |
| July | Ossian Sars | Echinodermata | Ophiurida | *Ophiopholis aculeata* | 7 | 0.002 |
| July | Ossian Sars | Echinodermata | Echinoida | *Strongylocentrotus pallidus* | 10 | 0.015 |
| July | Ossian Sars | Echinodermata | Camarodonta | NA | 19 | 0.452 |
| July | Ossian Sars | Mollusca | Chitonida | *Tonicella undocaerulea* | 224 | 0.108 |
| July | Ossian Sars | Mollusca | Adapedonta | *Hiatella arctica* | 899 | 0.205 |
| July | Ossian Sars | Mollusca | Myida | NA | 23 | 0.267 |
| July | Ossian Sars | Mollusca | Trochida | *Margarites helicinus* | 8 | 0.536 |
| July | Ossian Sars | Mollusca | Myoida | *Hiatella sp. K HML-2015* | 319 | 0.565 |
| July | Ossian Sars | Mollusca | Cardiida | NA | 26 | 0.706 |
| July | Ossian Sars | Ochrophyta | Ectocarpales | NA | 43 | 0.362 |
| July | Ossian Sars | Ochrophyta | Laminariales | NA | 2015 | 0.407 |
| July | Ossian Sars | Ochrophyta | Laminariales | *Halosiphon tomentosus* | 68 | 0.463 |
| July | Ossian Sars | Ochrophyta | Desmarestiales | NA | 34 | 0.675 |
