## Supplementary material for "Benthic diversity along an Arctic fjord: which are the key factors?": Table S2

Table S2: Paired Hutchinson tests' p-values for testing the significance of the difference between Shannon diversity of macroalgae and fauna for different periods (i.e., all periods combined, June, July, and August). Statistical significances are highlighted in gray (p-value < 0.5). Shannon diversity indexes are also indicated

### Macroalgae

| All periods | H' | Kongsbreen South | Ossian Sars | Bloomstrand East | Hansneset | French Bird Cliff | Kongsfjordneset |
|---|---|---|---|---|---|---|---|
| Kongsbreen South | 2.04 | NA | | | | | |
| Ossian Sars | 2.2 | 0.002 | NA | | | | |
| Bloomstrand East | 1.74 | 0.001 | 0.000 | NA | | | |
| Hansneset | 1.83 | 0.001 | 0.000 | 0.178 | NA | | |
| French Bird Cliff | 1.93 | 0.050 | 0.000 | 0.038 | 0.118 | NA | |
| Kongsfjordneset | 1.11 | 0.000 | 0.000 | 0.003 | 0.001 | 0.000 | NA |

| June | H' | Kongsbreen South | Ossian Sars | Bloomstrand East | Hansneset | French Bird Cliff | Kongsfjordneset |
|---|---|---|---|---|---|---|---|
| Kongsbreen South | 1.5 | NA | | | | | |
| Ossian Sars | 1.8 | 0.079 | NA | | | | |
| Bloomstrand East | 1.2 | 0.096 | 0.011 | NA | | | |
| Hansneset | NA | NA | NA | NA | NA | | |
| French Bird Cliff | 1.58 | 0.340 | 0.182 | 0.065 | NA | NA | |
| Kongsfjordneset | 1.1 | 0.005 | 0.000 | 0.278 | NA | 0.007 | NA |

| July | H' | Kongsbreen South | Ossian Sars | Bloomstrand East | Hansneset | French Bird Cliff | Kongsfjordneset |
|---|---|---|---|---|---|---|---|
| Kongsbreen South | 1.59 | NA | | | | | |
| Ossian Sars | 2.17 | 0.000 | NA | | | | |
| Bloomstrand East | 1.63 | 0.458 | 0.081 | NA | | | |
| Hansneset | 0.69 | 0.000 | 0.000 | 0.018 | NA | | |
| French Bird Cliff | 1.63 | 0.436 | 0.020 | 0.499 | 0.002 | NA | |
| Kongsfjordneset | 0.69 | 0.000 | 0.000 | 0.018 | NA | 0.002 | NA |

| August | H' | Kongsbreen South | Ossian Sars | Bloomstrand East | Hansneset | French Bird Cliff | Kongsfjordneset |
|---|---|---|---|---|---|---|---|
| Kongsbreen South | 1.3 | NA | | | | | |
| Ossian Sars | 1.76 | 0.057 | NA | | | | |
| Bloomstrand East | 1.36 | 0.388 | 0.050 | NA | | | |
| Hansneset | 1.72 | 0.057 | 0.441 | 0.041 | NA | | |
| French Bird Cliff | 1.43 | 0.344 | 0.179 | 0.402 | 0.196 | NA | |
| Kongsfjordneset | 1.08 | 0.165 | 0.007 | 0.042 | 0.003 | 0.133 | NA |

### Fauna

| All periods | H' | Kongsbreen South | Ossian Sars | Bloomstrand East | Hansneset | French Bird Cliff | Kongsfjordneset |
|---|---|---|---|---|---|---|---|
| Kongsbreen South | 2.76 | NA | | | | | |
| Ossian Sars | 3.18 | 0.023 | NA | | | | |
| Bloomstrand East | 2.53 | 0.229 | 0.016 | NA | | | |
| Hansneset | 2.59 | 0.209 | 0.000 | 0.408 | NA | | |
| French Bird Cliff | 2.55 | 0.230 | 0.008 | 0.467 | 0.437 | NA | |
| Kongsfjordneset | 2.17 | 0.090 | 0.018 | 0.216 | 0.150 | 0.189 | NA |

| June | H' | Kongsbreen South | Ossian Sars | Bloomstrand East | Hansneset | French Bird Cliff | Kongsfjordneset |
|---|---|---|---|---|---|---|---|
| Kongsbreen South | 2.38 | NA | | | | | |
| Ossian Sars | 2.95 | 0.011 | NA | | | | |
| Bloomstrand East | 2.53 | 0.263 | 0.002 | NA | | | |
| Hansneset | 1.3 | 0.007 | 0.001 | 0.005 | NA | | |
| French Bird Cliff | 2.61 | 0.153 | 0.007 | 0.237 | 0.004 | NA | |
| Kongsfjordneset | 1.62 | 0.005 | 0.000 | 0.001 | 0.191 | 0.001 | NA |

| July | H' | Kongsbreen South | Ossian Sars | Bloomstrand East | Hansneset | French Bird Cliff | Kongsfjordneset |
|---|---|---|---|---|---|---|---|
| Kongsbreen South | 2.66 | NA | | | | | |
| Ossian Sars | 2.95 | 0.011 | NA | | | | |
| Bloomstrand East | 1.88 | 0.002 | 0.000 | NA | | | |
| Hansneset | 2.05 | 0.001 | 0.000 | 0.247 | NA | | |
| French Bird Cliff | 2.29 | 0.014 | 0.000 | 0.049 | 0.093 | NA | |
| Kongsfjordneset | 1 | 0.001 | 0.001 | 0.009 | 0.004 | 0.002 | NA |

| August | H' | Kongsbreen South | Ossian Sars | Bloomstrand East | Hansneset | French Bird Cliff | Kongsfjordneset |
|---|---|---|---|---|---|---|---|
| Kongsbreen South | 1.69 | NA | | | | | |
| Ossian Sars | 2.35 | 0.006 | NA | | | | |
| Bloomstrand East | 1.87 | 0.343 | 0.139 | NA | | | |
| Hansneset | 2.44 | 0.003 | 0.282 | 0.103 | NA | | |
| French Bird Cliff | 1.59 | 0.325 | 0.000 | 0.258 | 0.000 | NA | |
| Kongsfjordneset | 2.12 | 0.105 | 0.225 | 0.300 | 0.148 | 0.050 | NA |
