## Supplementary material for "Benthic diversity along an Arctic fjord: which are the key factors?": Figure S1

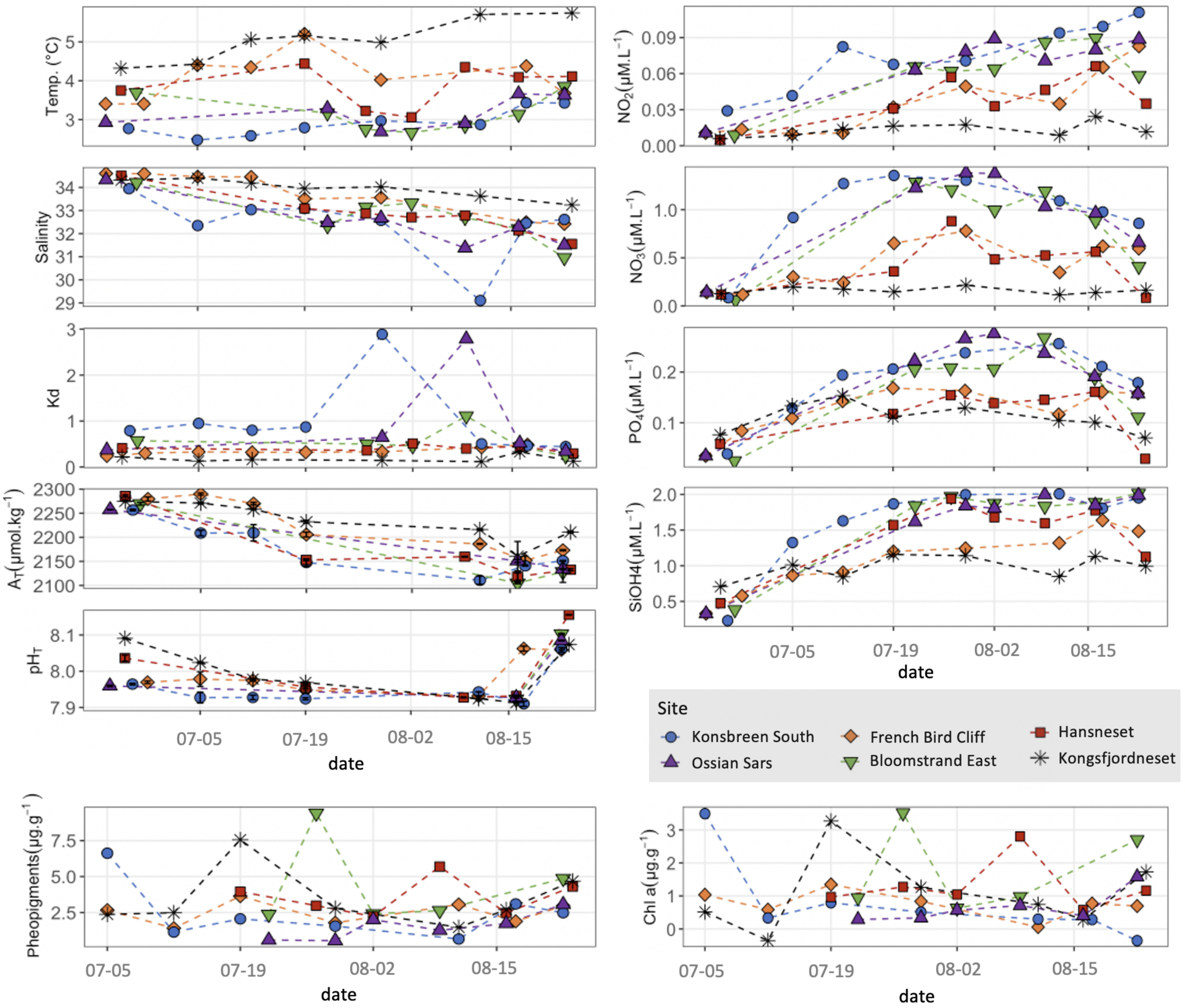


Figure S1: Environmental parameters measurements at each site within the summer 2021 in Kongsfjorden
